# Variability in the Analgesic Response to Ibuprofen Following Third-Molar Extraction is Associated with Differences in Activation of the Cyclooxygenase Pathway

**DOI:** 10.1101/467407

**Authors:** Katherine N. Theken, Elliot V. Hersh, Nicholas F. Lahens, Hyo Min Lee, Eric J. Granquist, Helen Giannakopoulos, Lawrence M. Levin, Stacey A. Secreto-Dankanich, Gregory R. Grant, John A. Detre, Garret A. FitzGerald, Tilo Grosser, John T. Farrar

**Affiliations:** University of Pennsylvania Perelman School of Medicine, Institute for Translational Medicine and Therapeutics; University of Pennsylvania School of Dental Medicine, Oral Surgery and Pharmacology; McGill University, Montreal Neurological Institute and Hospital, Department of Neurology and Neurosurgery; University of Pennsylvania Perelman School of Medicine, Neurology; University of Pennsylvania Perelman School of Medicine, Biostatistics and Epidemiology

## Abstract

The analgesic efficacy of non-steroidal anti-inflammatory drugs (NSAIDs) has long been recognized to be limited by substantial interindividual variability in pain relief, but the underlying mechanisms are not well understood. We performed pain phenotyping, functional neuroimaging, pharmacokinetic/pharmacodynamic assessments, inflammation biomarkers, and gene expression profiling in healthy subjects who underwent surgical extraction of bony impacted third molars, in order to characterize factors associated with heterogeneity in response to ibuprofen.

Subjects were treated with rapid-acting ibuprofen (400 mg; N=19) or placebo (N=10) in a randomized, double-blind design. Compared to placebo, ibuprofen-treated subjects exhibited greater reduction in pain scores, alterations in regional cerebral blood flow in brain regions associated with pain processing, and inhibition of ex vivo cyclooxygenase activity and urinary prostaglandin metabolite excretion as indices of biochemical drug action (p<0.05). As expected, ibuprofen-treated subjects could be stratified into partial responders (N=9, required rescue medication within the dosing interval) and complete responders (N=10, no rescue medication). This was also reflected by differences in pain scores (p<0.01) as early as 30 minutes following drug administration (p<0.05). Variability in analgesic efficacy was not associated with demographic or clinical characteristics, ibuprofen pharmacokinetics, metabolizing enzyme genotype, or the degree of cyclooxygenase inhibition by ibuprofen. However, complete responders had higher concentrations of inflammatory biomarkers in urine and serum, than partial responders. Specifically, a stable urinary prostaglandin E_2_ metabolite, serum TNFα and IL-8 were higher in patients who did not require rescue medication compared those who did (p < 0.05). RNAseq gene expression analysis in PBMCs collected after surgery and ibuprofen administration showed enrichment of inflammation related pathways among genes differentially expressed (q < 0.2) between complete and partial responders

These findings suggest that patients who receive substantial pain relief from ibuprofen have a more pronounced activation of the prostanoid biosynthetic pathway and regulation of the inflammatory pain phenotype differs from those patients who are insufficiently treated with ibuprofen alone and may require an opioid or other therapeutic intervention.

## Introduction

Although acute pain resulting from injury or other tissue damage can serve an adaptive function by promoting behaviors that limit the chance of further trauma, inadequate pain management in the post-operative setting can delay healing and adversely affect mental well-being. Opioid analgesics are an important component of post-operative pain management in many patients. However, over-prescription of opioids for surgical pain, typically 2-5 times more than patients actually use, has contributed to the opioid epidemic [1, 2]. Thus, there is a need to consider alternative therapeutic options for those patients, whose pain can be appropriately managed with non-addictive analgesics, including nonsteroidal anti-inflammatory drugs (NSAIDs).

NSAIDs exert their pharmacologic effects via inhibition of one or both cyclooxygenase (COX) enzymes (COX-1 and COX-2), which catalyze the first committed step in the synthesis of prostanoids (prostaglandin (PG) E_2_, prostacyclin (PGI_2_), PGD_2_, PGF_2α_, and thromboxane (Tx) A_2_) from arachidonic acid [3, 4]. In the setting of inflammatory pain, these lipid mediators, particularly PGE_2_ and PGI_2_, act locally on their respective G protein coupled receptors to promote peripheral [5-9] and central [10, 11] sensitization, thus rendering the nociceptive system more excitable. Because NSAIDs inhibit the formation of prostanoids that promote peripheral and central sensitization, rather than directly modulating the nociceptive system, they are effective in treating pain in which activation of the COX pathway is a key mechanism.

One such situation is surgical extraction of third molars, a procedure undergone by approximately 5 million patients per year in the United States [12, 13]. The soft tissue and bony trauma associated with third molar extraction surgery liberates key inflammatory mediators, including prostanoids, activating and sensitizing free nerve endings at the surgical site [6, 14]. At the population level, standard NSAID doses are on average at least as effective as standard opioid doses following third molar extraction surgery [15-18]. However, there is considerable variability in the analgesic response to NSAIDs at the individual level, with 20-30% of patients requiring opioid rescue medication within 6 hours of the initial NSAID dose [19, 20]. In order to avoid undertreating patients who will not respond adequately to NSAIDs, oral surgeons routinely prescribe opioids to be taken if needed, resulting in unnecessary prescriptions for a majority of patients who do not require them or only require minimal dosing [19, 21].

The application of precision medicine approaches to pain management after third molar extraction would facilitate the optimization of NSAID therapy and limit opioid prescriptions to those patients who would not attain adequate pain relief with NSAIDs. However, the development of such approaches requires a much better understanding of the molecular mechanisms that contribute to variation in analgesic efficacy of NSAIDs. Therefore, we performed a deep phenotyping study incorporating functional neuroimaging, pharmacokinetic/pharmacodynamic assessments, biochemical assays, and gene expression analysis to characterize the factors that are associated with inter-individual variability in analgesic efficacy of ibuprofen following third molar extraction surgery.

## Methods

### Subjects

Healthy subjects (≥18 years of age) were recruited from patients referred to the Oral and Maxillofacial Surgery Service at the School of Dental Medicine and the Hospital of the University of Pennsylvania for extraction of one or more partially or fully bony impacted third molars. Upper third molars could be included in the surgical plan, if appropriate, but at least one lower tooth was required because upper extractions alone do not consistently lead to a significantly painful post-operative state [22]. Subjects were excluded if they had 1) pain lasting more than one day for the two weeks prior to surgery, 2) a positive pregnancy test, 3) any serious medical illness that would make participation unsafe, 4) any history of psychological illness requiring medication or other treatment for more than three months, 5) any history of claustrophobia that could affect the subject’s ability to tolerate the MRI study, 6) bleeding disorder or current use of anticoagulants by patient history, 7) use of any opioid medication more than three times in the last week, or 8) the presence of any implanted devices or metal objects in their body that might exclude them from having a MRI.

### Study procedures

The study protocol was approved by the University of Pennsylvania Institutional Review Board, and all subjects provided informed consent. One week prior to their scheduled surgery, subjects attended a study visit to demonstrate the imaging process and to fill out measures for demographics, personality, mood, and pain coping skills, as previously described [23]. They were asked to abstain from analgesics, including products containing NSAIDs, aspirin, and acetaminophen, high dose vitamins, and nutritional supplements until after surgery.

The study procedures are summarized in Figure 1. On the day of surgery, baseline blood and spot urine samples were collected. Subjects then underwent third molar extraction with 3% mepivacaine plain and 2% lidocaine plus 1:100,000 epinephrine for local anesthesia and nitrous oxide/oxygen and midazolam titrated to effect for sedation to ensure adequate pain management during the procedure and relatively rapid dissipation of the effect once surgery was complete. A trauma score was determined based on the difficulty of extraction, as previously described [24]. After surgery subjects reported pain intensity every 15 minutes using the 0-10 Numeric Rating Scale (NRS-PI), where 0=no pain and 10=worst pain imaginable.

**Figure 1:**
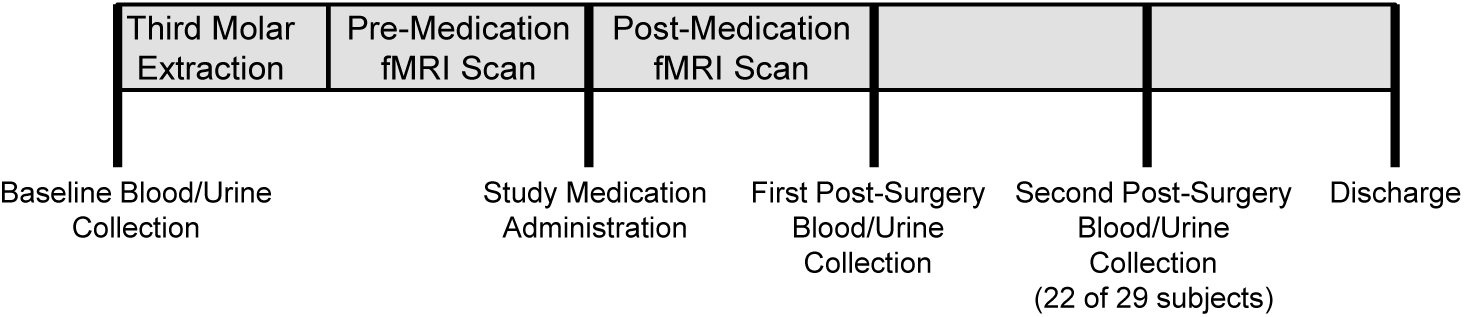
Study design

Approximately 45 minutes after completion of surgery, subjects were placed in the MRI to begin functional imaging data collection in intervals of 15 minutes. When the study subjects requested medication and reported a pain score ≥4/10 or indicated that their pain was no longer tolerable, they received a dose of rapid-acting ibuprofen sodium (Advil^®^ filmtabs) or matching placebo by mouth according to their randomization assignment. After administering the study medication, MRI scanning continued for up to 60 minutes. For both the pre-medication and post-medication scans, the imaging data was collected with 5 minutes of registrations scans, followed by continuous functional scanning in 15-minute intervals with pain levels recorded between each scanning segment. Subjects were encouraged to allow at least 60 minutes for the study medication to take effect before deciding if it had been ineffective, but rescue medication (hydrocodone 5 mg/ acetaminophen 325 mg) was allowed any time upon request.

Immediately after the scanning session, post-surgery blood and spot urine samples were collected, and subjects were returned to the post-operative observation area for continued pain assessment every 30 minutes for a total of two hours after treatment. After the first 7 subjects were enrolled, the study procedure was amended to include a second post-surgery blood and urine sample collection approximately 3 hours after study drug administration. Once medically stable, the subjects were discharged home with a prescription for a 3-day supply of hydrocodone 5 mg/ acetaminophen 325 mg.

### fMRI Procedure

A 3 Tesla Siemens Trio MRI system with a 32-channel head receiver was used. Subjects received ear plugs to reduce scanner noise, and foam padding was used to limit head motion. The scanning protocol began with a 3-axis localizer (1 min) followed by a 3-dimensional MPRAGE anatomical scan obtained with 1 mm isotropic resolution (5 min) used to define brain anatomy. The remainder of the scanning consisted of sequential measurements of regional cerebral blood flow (CBF) using arterial spin labeling (ASL)-MRI obtained with pulsed continuous labeling [25, 26]. Each scan consisted of 30 label/control pairs for a scan duration of 4 minutes per period.

Perfusion fMRI data was analyzed offline using established procedures by an investigator blinded to treatment assignment. Raw perfusion data was cleaned using motion correction and outlier elimination [27], and mean CBF maps for each time point were calculated and coregistered into a standard anatomical space using the PET module of SPM12 and the MPRAGE anatomical data. The time course of CBF changes in *a priori* regions of interest were extracted using the PICKATLAS [28] utility within SPM and the NeuroSynth pain (http://neurosynth.org/). The extracted CBF data from regions of interest were evaluated with standard statistical analyses to test for effects of rapid-acting ibuprofen sodium vs. placebo on neural activity. Whole-brain topologic inference was applied via the false discovery rate (FDR) method based on Gaussian random field theory, however, due to the small groups sizes no uncorrected p value survived FDR estimation. Uncorrected p values were displayed to illustrate trends using color gradation based on Cohen’s effect size, only in the areas that survived p < 0.05 (uncorrected) thresholding.

### Quantification of COX activity and plasma drug concentrations

COX-1 activity *ex vivo* was evaluated by quantifying serum thromboxane B_2_ levels, as previously described [29]. Briefly, whole blood was collected into vacuum tubes containing clot activator and incubated in a water bath at 37°C for 1 hour. Serum was separated by centrifugation and stored at −80°C until analysis by liquid chromatography/mass spectrometry (LC-MS).

COX-2 activity *ex vivo* was evaluated by quantifying plasma PGE_2_ levels following lipopolysaccharide (LPS) stimulation in whole blood, as previously described [30]. Briefly, heparinized whole blood was treated with aspirin (1mM) and incubated at room temperature for 15 minutes. LPS (E. coli, serotype O111:B4, 10 μg/ml whole blood) was added, and the sample was incubated in a water bath at 37°C for 24 hours. Plasma was separated by centrifugation and stored at −80°C until analysis by LC-MS.

COX activity *in vivo* was determined by quantification of urinary prostanoid metabolites by liquid chromatography mass spectrometry (LC-MS) as previously described [31]. Systemic production of PGI_2_, PGE_2_, PGD_2_, and thromboxane (Tx) A_2_ was determined by quantifying their major urinary metabolites: 2,3-dinor 6-keto-PGF_1α_ (PGIM), 7-hydroxy-5,11-diketotetranorprostane-1,16-dioic acid (PGEM), 11,15-dioxo-9α-hydroxy-2,3,4,5-tetranorprostan-1,20-dioic acid (PGDM), and 2,3-dinor TxB_2_ (TxM), respectively. Results were normalized to urinary creatinine.

Plasma concentrations of ibuprofen and acetaminophen were quantified by LC-MS as previously described [32].

### Serum cytokines

Concentrations of interleukin (IL)-6, IL-1β, IL-8, IL-10, tumor necrosis factor (TNF)-α, and monocyte chemoattractant protein (MCP)-1 in serum were quantified by MILLIPLEX multiplex assay (Millipore) by the Radioimmunoassay and Biomarkers Core at the Diabetes Research Center at the University of Pennsylvania. The levels of IL-1β were below the limit of detection in the majority of samples, so further statistical analysis was not performed for this analyte.

### CYP2C9 Genotyping

Genomic DNA was isolated from buffy coat samples using the PureLink Genomic DNA Mini Kit (Invitrogen). TaqMan SNP Genotyping assays (ThermoFisher) were used to genotype *CYP2C9*\**2* (C_25625805_10) and *CYP2C9*\**3* (C_2710892_10) per the manufacturer’s instructions.

### Gene expression analysis

Peripheral blood mononuclear cells (PBMCs) were isolated from whole blood using a standard Ficoll-Paque density gradient centrifugation method. Total RNA was isolated using the RNeasy Miniprep Kit (QIAGEN) per the manufacturer’s instructions. Total RNA from each sample was converted to sequencing libraries using the SMARTer Stranded Total RNA-Seq Kit v2 - Pico Input Mammalian (Clontech), following the manufacturer’s protocol. Each sample was prepared using a unique dual-barcode combination. All libraries were pooled together at equimolar concentrations and test-sequenced with a single MiSeq run (2×150bp reads). The results of the MiSeq run were used to assess the balance of reads generated from each library in the pool, and the relative concentrations of the libraries in the pool were adjusted accordingly. The re-balanced pool was then sequenced across 6 lanes on a HiSeq 4000 (2×150bp reads). Following QC of the pooled HiSeq data, a second pool of the libraries was sequenced to increase read depth.

Illumina adapter sequences were trimmed from the raw, gzipped FASTQ files using the BBDuk tool from the BBTools suite v37.99 (https://jgi.doe.gov/data-and-tools/bbtools/). Reads from the trimmed FASTQ files were aligned to GRCh38 build of the human reference genome using STAR v2.6.0c [33] and gene models from v92 of the Ensembl annotation [34].

Data were normalized and quantified using the Pipeline Of RNA-seq Transformations v0.8.5b-beta (PORT; https://github.com/itmat/Normalization), a resampling-based method that accounts for confounding factors like read depth, ribosomal RNA, mitochondrial RNA content, and genes with extremely high expression. PORT was run at the gene level in strand-specific mode and was provided with gene models from v92 of the Ensembl annotation. During normalization, 11 of a total of 77 samples were excluded for low read depth. Following normalization, pairwise differential expression (DE) analyses were performed on the gene-level read counts using *voom*-*limma* software package v3.34.0 [36]. The data are accessible through GEO Series accession number GSE120596. Only genes with >0 reads across all samples in at least one of the two compared groups were used for pairwise DE analyses. Pathways enriched in each of the DE gene lists were identified using Ingenuity Pathway Analysis (IPA; Qiagen) [37].

### Statistical analysis

Data are reported as mean ± standard deviation or median (25^th^ percentile, 75^th^ percentile). Baseline characteristics and biochemical measurements were compared by t-test or ANOVA or their non-parametric equivalents, as appropriate. Time to rescue medication treatment was evaluated by log-rank test. Post-surgery measurements of COX activity *ex vivo*, urinary PG metabolite levels, and serum inflammatory mediators were normalized to baseline values for each subject and compared at each time point by Wilcoxon rank-sum test. P<0.05 was considered statistically significant.

## Results

### Study cohort

The study cohort included 29 healthy adults (16 men, 13 women) with a mean age of 24.9±3.59 years. Ten subjects received placebo, and 19 subjects received ibuprofen. Usable fMRI scans were obtained in 24 subjects (placebo: N=6; ibuprofen: N=18), others had machine or procedural issues or excess movement artifact sufficient to prevent appropriate analysis. The median maximum pain score before study drug administration (0-10 scale) was 7 (5, 8) which was administered with a mean time of 1.85±0.55 h after surgery. The first post-surgery and second post-surgery sample collections occurred 1.51±0.47 h and 3.17±0.42 h after study drug administration, respectively.

### Activity of ibuprofen

Ibuprofen was significantly more effective than placebo in relieving pain following third molar extraction. Thus, the median pain intensity difference (PID; maximum pain score before study drug – minimum pain score before rescue medication treatment) was 3 (2, 5) in the ibuprofen group, compared to −0.5 (−2, 1.25) in the placebo group (p<0.001). The patients’ global assessment of pain relief also favored ibuprofen with 16 of 19 subjects who received ibuprofen rating their pain as “much better” or “very much better” after study drug treatment, compared to 0 of 10 subjects who received placebo (p<0.0001; Fisher exact test).

The onset of pain relief was detectable between 15 and 30 minutes following drug administration as indicated by a difference in the slope of pain intensity scores between these time points (p<0.001 placebo vs. ibuprofen). Functional neuroimaging analysis was restricted to these early time points (predrug = 0 min, 15 min and 30 min), because no patient in the placebo group was able to remain in the scanner longer than 30 min following placebo administration, while some patients in the ibuprofen group tolerated 60 min to 75 min of post drug scan time. Additionally, poor image quality primarily due to motion artifacts precluded analysis in 4 out of 10 placebo group patients, but only 1 out of 19 ibuprofen group patients. Despite these limitations a significant change in CBF indicative of the analgesic effect of ibuprofen was detectable between 15 and 30 minutes in the summary analysis of the brain’s pain processing regions (Supplemental Figure 1). This was primarily driven by perfusion changes in the insula, the anterior cingulate cortex and the secondary somatosensory cortex.

Ibuprofen inhibited COX activity, as indicated by *ex vivo* whole blood assays and quantification of urinary PG metabolites (Supplemental Figure 2). COX-1 activity *ex vivo* was inhibited by >90% at both post-surgery time points in ibuprofen-treated subjects (p<0.001). COX-2 activity *ex vivo* was also inhibited in ibuprofen-treated subjects at the first post-surgery time point (ibuprofen: 25.3±21.0% of baseline; N=17 vs. placebo: 40.0±19.3% of baseline; N=10; p=0.0404). Some subjects had received rescue medication prior to the first post-surgery sample collection, and acetaminophen, which was a component of the rescue medication (hydrocodone 5 mg/ acetaminophen 325 mg), may also inhibit COX-2 activity [38]. When the comparison was restricted to subjects with plasma acetaminophen concentrations below the limit of quantitation, the inhibition of COX-2 activity *ex vivo* at the first post-surgery time point was also apparent (ibuprofen: 24.7±12.4% of baseline; N=13 vs. placebo: 51.3±12.6% of baseline; N=5; p=0.003). Ibuprofen-treated subjects also exhibited significantly lower levels of the urinary metabolites of PGE_2_, PGI_2_, and PGD_2_ relative to baseline at the second post-surgery time point, compared to the placebo group (p<0.001).

### Variability in the ibuprofen response

The time to rescue medication treatment is shown in Figure 2A. All ten subjects in the placebo group requested opioid rescue medication, with a median time to rescue of 53 (48, 55) minutes after study drug administration. Nine subjects in the ibuprofen group requested opioid rescue medication and were categorized as “partial responders”. The time to rescue medication treatment was significantly longer in partial responders compared to the placebo group (p<0.001; log-rank test), with a median time to rescue of 105 (82, 133) minutes after study drug administration. The remaining ten subjects in the ibuprofen group did not require additional analgesic medication 4 hours after dosing and were categorized as “complete responders”. Pain intensity scores after study medication administration also differed among the response groups (Figure 2B). The median PID was 5 (3, 6.25) in complete responders, compared to 2 (1, 3.5) in partial responders and −0.5 (−2, 1.25) in placebo-treated subjects (p=0.0003, Kruskal-Wallis test). The separation into the three groups was detectable as early as 30 minutes following study drug administration based on reported pain scores (Figure 2C). Functional neuroimaging did not allow distinction between complete and partial responders, within 30 minutes of drug administration and the number of patients who did not tolerate prolonged scan time precluded sufficiently powered analysis at later time points (Figure 2C and D).

**Figure 2:**
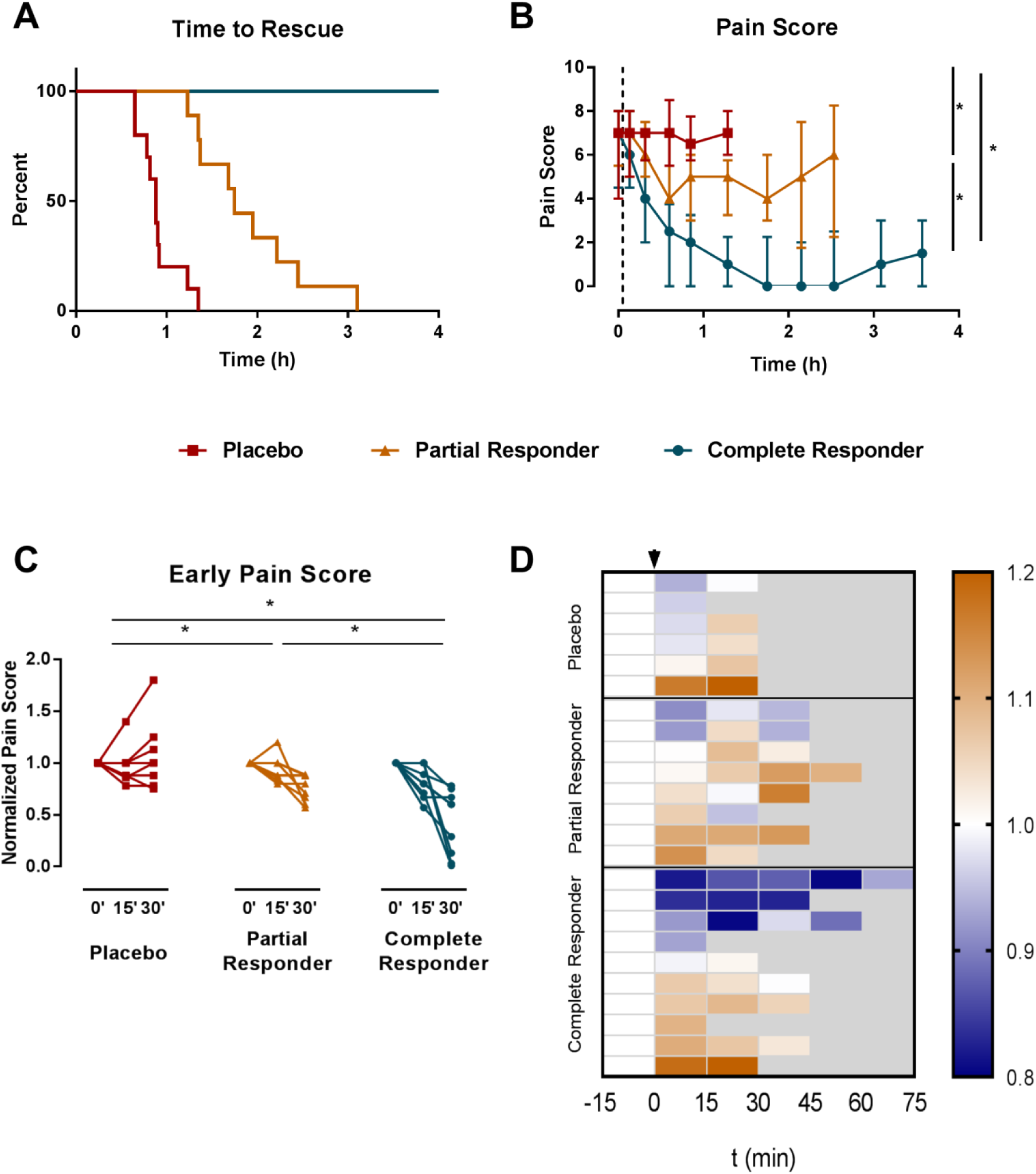
Inter-individual variability in analgesic response to ibuprofen. **A)** Kaplan-Meier curves depicting time to rescue medication administration by response group (placebo: n=10; partial responders: n=9; complete responders: n=10; p<0.001 for all comparisons in the log-rank test). **B)** Pain scores at each pain assessment prior to rescue medication administration by response group (*p<0.01; Kruskal-Wallis test). Vertical dotted line indicates time of study drug administration. **C)** Change of pain scores relative to pre-drug (0 min) scores up to 30 minutes (*p<0.01; Kruskal-Wallis test). **D)** Heatmap depicting change from pre-drug of the integrated CBF measurements in pain regions (NeuroSynth pain map) by individual patients.

### Variability in the inflammatory response to surgery

No significant differences in demographic or clinical characteristics (e.g. number of extracted teeth, trauma score, etc) were observed between complete and partial responders (Table 1), and baseline COX activity *ex vivo* and levels of PG metabolites did not differ among the response groups (Supplemental Table 1). Similar ibuprofen plasma concentrations and degree of inhibition of COX activity *ex vivo* were observed in complete and partial responders at both post-surgery time points (Supplemental Figure 3), suggesting that differences in the ibuprofen response cannot be explained by differences in pharmacokinetics. There was no significant difference in the frequency of *CYP2C9*\**2* or *CYP2C9*\**3* variant alleles between the response groups (data not shown).

**Table 1:** Baseline characteristics by response group

|  | Placebo | Partial Responder | Complete Responder |
| --- | --- | --- | --- |
| N | 10 | 9 | 10 |
| Men/Women | 5/5 | 4/5 | 7/3 |
| Age (y) | 26.1±3.9 | 25.7±2.9 | 23.1±3.4 |
| Race |  |  |  |
| White | 1 | 3 | 5 |
| Asian | 5 | 5 | 2 |
| African-American | 4 | 0 | 0 |
| Other | 0 | 1 | 3 |
| Length of surgery (min) | 40.0±15.0 | 36.6±21.4 | 38.6±32.2 |
| Number of teeth | 4 (3,4) | 4 (4,4) | 3.5 (2.25,4) |
| Trauma Score | 8 (6.25,8) | 7 (6,8) | 7 (5.25,7.75) |
| Max Pain Score | 7 (5, 8) | 7 (6,8) | 7 (5,7.75) |

Interestingly, partial responders exhibited greater suppression of urinary PGEM (partial responders: 45.3±14.9% of baseline vs. complete responders: 75.7±23.7% of baseline; p=0.0021), PGIM (partial responders: 70.6±43.1% of baseline vs. complete responders: 126.5±43.7% of baseline; p=0.0133), and tended that way for PGDM (partial responders: 66.5±20.7% of baseline vs. complete responders: 83.1±12.6% of baseline; p=0.0947) at the first post-surgery time point, compared to complete responders (Figure 3). Greater suppression of urinary PG metabolites was observed at the second post-surgery time point. The difference between partial and complete responders persisted for PGEM (partial responders: 21.2±7.8% of baseline vs. complete responders: 38.4±11.9% of baseline; p=0.0082), but PGIM and PGDM were suppressed to a similar degree (∼50% of baseline) in both partial and complete responders. At the second post-surgery time point, serum TNF-α (partial responders: 73.1±26.8% of baseline vs. complete responders: 186.2±143.3% of baseline; p=0.0127) and IL-8 (partial responders: 73.5±40.6% of baseline vs. complete responders: 199.4±88.1% of baseline; p=0.0027) were induced to a greater extent in complete responders than in partial responders, while no significant differences between the groups were observed for IL-6, IL-10, and MCP-1 (Figure 4). No significant differences in serum levels of these inflammatory mediators were observed at baseline (Supplemental Table 2).

**Figure 3:**
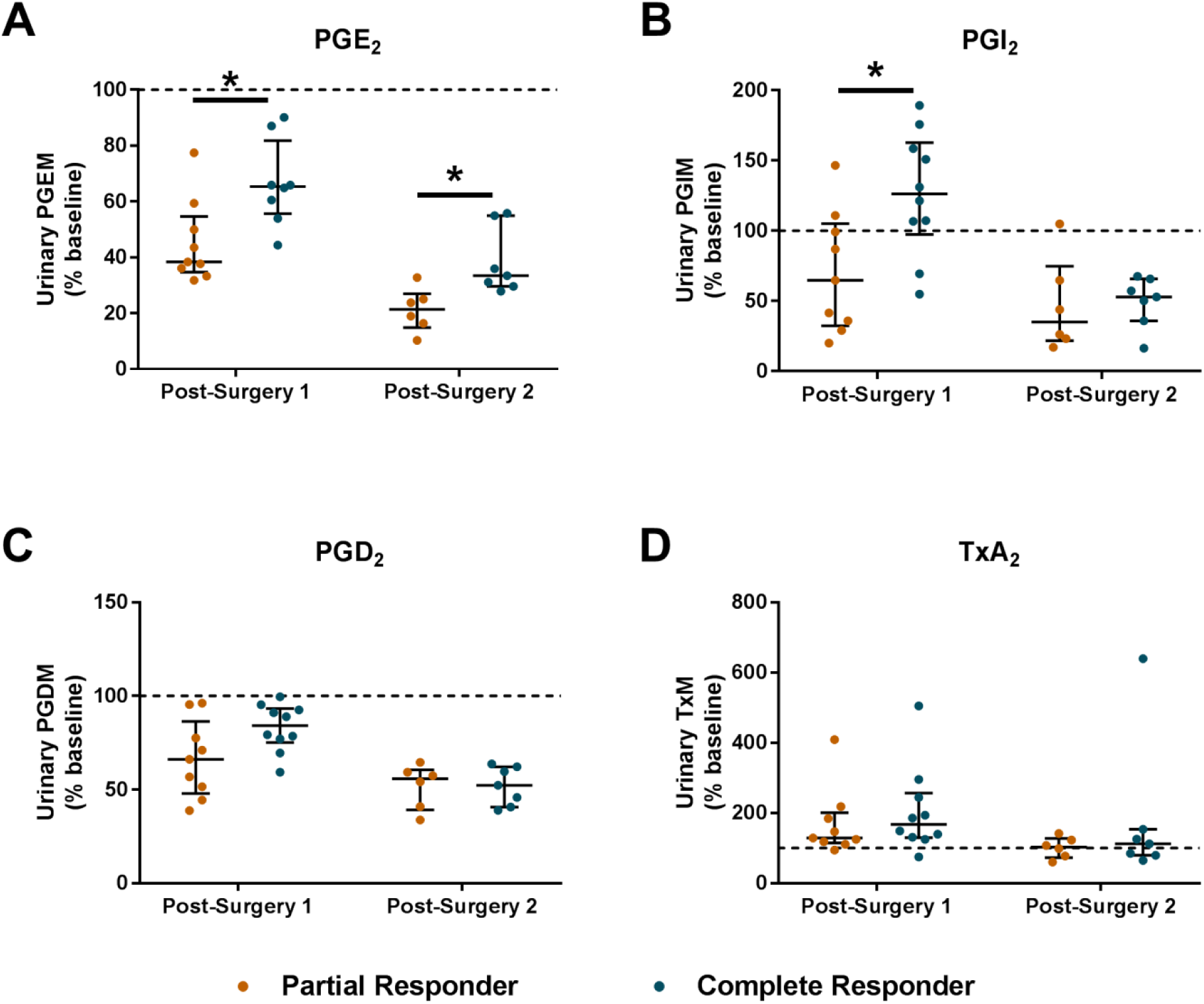
Comparison of urinary PG metabolite levels after surgery in partial (gold) and complete (teal) responders. *p<0.05 by Wilcoxon rank-sum test.

**Figure 4:**
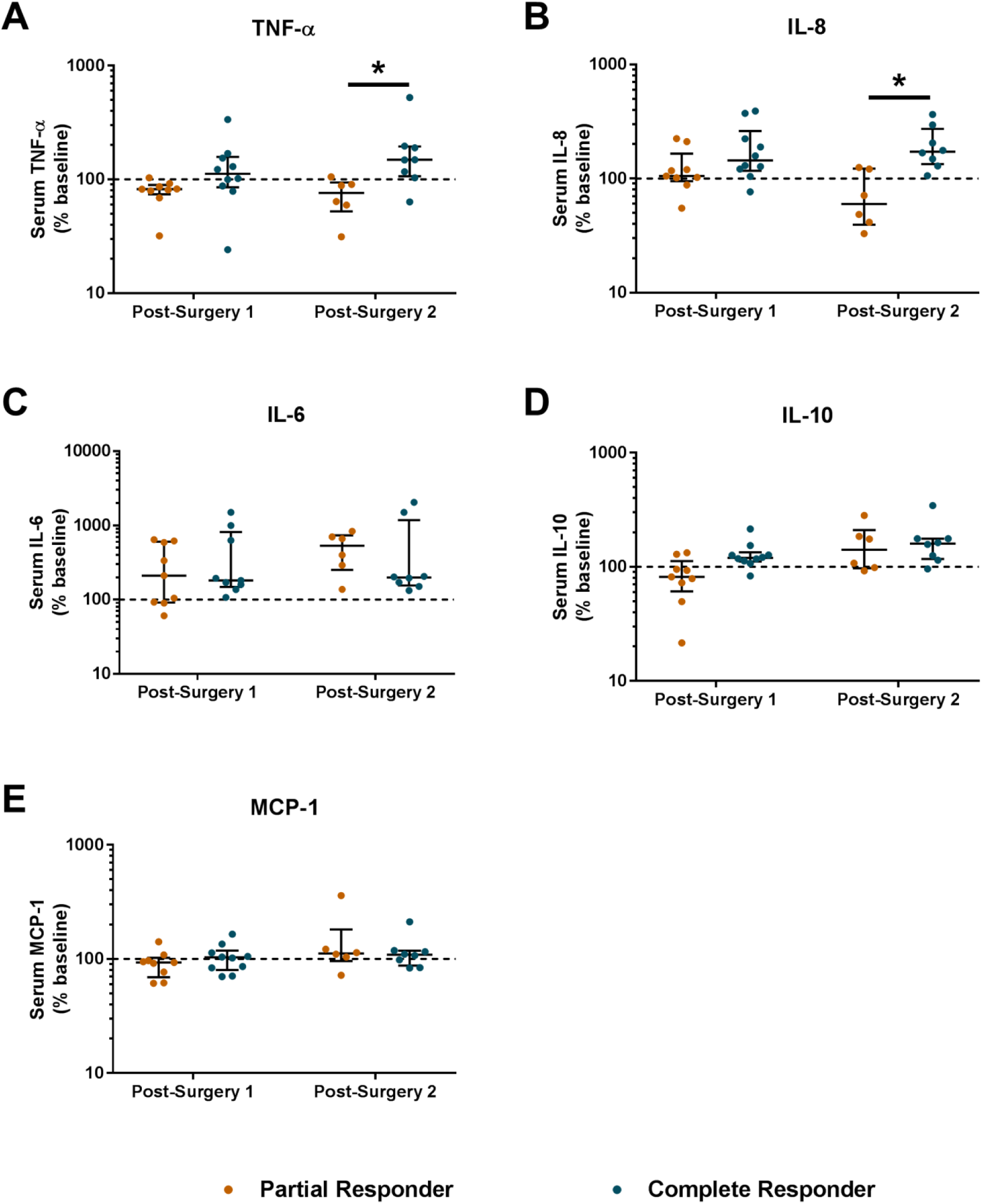
Comparison of serum inflammatory mediators after surgery in partial and complete responders. *p<0.05 by Wilcoxon rank-sum test.

Gene expression in PBMCs did not differ among the response groups at baseline. After surgery and study drug treatment, the placebo group had the most changes in gene expression, with 431 genes at the first post-surgery time point and 2653 genes at the second post-surgery time point differentially expressed compared to baseline (q<0.05). Partial responders exhibited fewer changes in gene expression, with 99 differentially expressed genes (DEGs) at the first post-surgery time point and 1092 DEGs at the second post-surgery time point (q<0.05). Very few genes were differentially expressed compared to baseline in complete responders, with 7 DEGs at the first post-surgery time point and 47 DEGs at the second post-surgery time point (q<0.05). Subsequent analyses focused on the second post-surgery time point because the majority of DEGs observed at the first post-surgery time point within each group were also differentially expressed at the second post-surgery time point (data not shown). There was significant overlap in the DEGs among the three groups (Figure 5A). Pathway analysis of the DEGs indicated that these were enriched for pathways related to inflammation, but specific pathways differed among the ibuprofen response groups (Supplemental Figures 4-6). At the second post-surgery time point, 1345 genes were differentially expressed between partial and complete responders (q<0.2; Figure 5B), with enrichment for inflammatory pathways (Table 2). Expression plots for select genes from these pathways are shown in Supplemental Figure 7. For the majority of these inflammatory genes, partial responders had significantly higher expression compared to complete responders.

**Table 2:** Pathway analysis of genes differentially expressed between partial and complete responders at the second post-surgery time point (q<0.2).

| Pathway Name | p-value |
| --- | --- |
| Fc Receptor-mediated Phagocytosis in Macrophages and Monocytes | $5.31 \times 10^{-6}$ |
| TREM1 Signaling | $5.98 \times 10^{-6}$ |
| Neuroinflammation Signaling Pathway | $3.62 \times 10^{-5}$ |
| CD28 Signaling in T Helper Cells | $5.27 \times 10^{-5}$ |
| Production of Nitric Oxide and Reactive Oxygen Species in Macrophages | $7.32 \times 10^{-5}$ |

**Figure 5:**
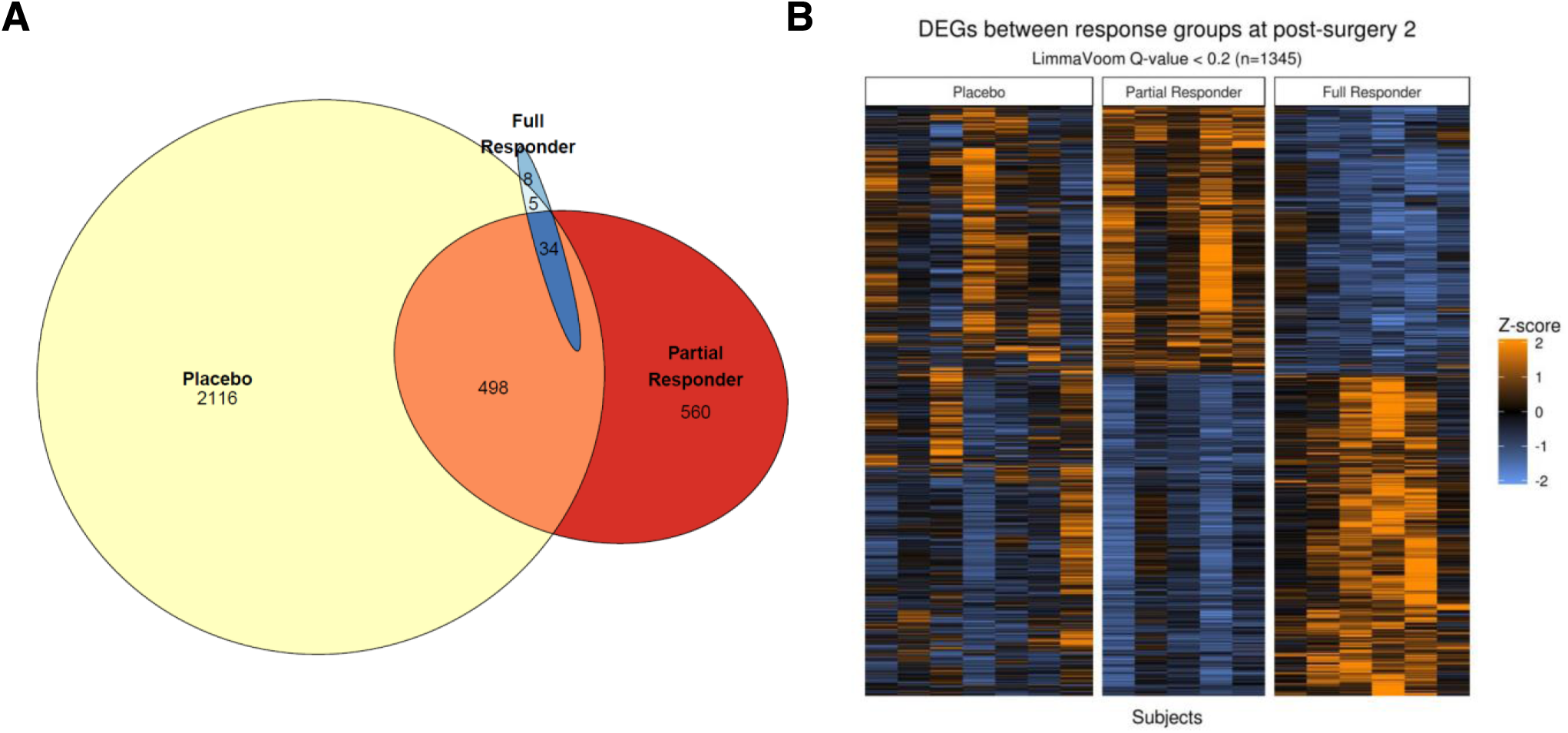
RNA sequencing analysis of PBMCs. **A)** Venn diagram depicting overlap among the response groups in genes differentially expressed at the second post-surgery time point compared to baseline (q<0.05). **B)** Heatmap depicting differentially expressed genes (DEGs) between partial and complete responders at the second post-surgery time point (q<0.2).

## Discussion

Pain is a complex multidimensional experience that reflects the interaction between nociceptive, affective, and cognitive processes [39, 40]. Given the diverse mechanisms that contribute to pain, it has long been recognized that there is substantial inter-individual variability in the effectiveness of all analgesics, including NSAIDs [16, 19, 20, 41]. For example, it has been estimated that only half of arthritis patients prescribed NSAIDs will have a moderate or better pain relief response [41]. Studies in acute post-surgical pain following third molar extraction have demonstrated that, while NSAIDs are highly effective on average, 20-30% of patients required opioid rescue medication within 6 hours of the initial NSAID dose indicating that these were individuals in whom NSAIDs failed to provide sufficient pain relief throughout the dosing interval [19, 20]. Currently, pain therapy is configured on a trial-and-error approach, often involving several iterations of switching drugs and adjusting doses. Applying precision medicine approaches to the personalization of pain therapy promises to accelerate treatment optimization for individual patients. However, the development of such treatment algorithms is limited by our lack of understanding of the mechanisms underlying inter-individual variability in analgesic efficacy. Here we demonstrate that variability in the response to ibuprofen following third molar extraction is detectable across multiple diagnostic domains - behavioral, brain imaging, and markers of systemic inflammation - indicating that partial and complete analgesic responses to ibuprofen reflect internally consistent phenotypes. In addition, we find that activation of the prostanoid biosynthetic pathway associated with surgical trauma differs between complete and partial responders, suggesting that the response phenotype relates to the mechanism of drug action.

Thus, a key strength of our study is the application of an array of complementary techniques to differentiate between partial and complete responders. Pain and analgesic efficacy are challenging to quantify in a clinical setting due to the inherent subjectivity in the experience of pain and imprecision of pain rating scales [42]. Functional neuroimaging has expanded the understanding of the neural basis of pain mechanisms and may provide an objective biomarker of efficacy of pain treatment. A prior study that used ASL-fMRI to quantify the effect of ibuprofen administration after third molar extraction demonstrated that the analgesic effect of ibuprofen was associated with decreases in CBF in brain regions known to be involved in the perception of post-surgical pain [43]. We observed similar results in our ibuprofen-treated subjects. Moreover, these alterations in CBF were most apparent in complete responders who also reported a greater improvement in pain scores.

Notably, response to ibuprofen in our cohort could not be predicted based on clinical characteristics, ibuprofen pharmacokinetics, or pharmacodynamics. Rather, we observed that complete responders exhibited higher levels of urinary PG metabolites at the first post-surgery time point, compared to partial responders. Because the concentrations of PG metabolites in urine reflect COX activity *in vivo* over the entire collection interval, this measurement is influenced both by the degree of activation of the COX pathway in response to third molar extraction surgery, as well as inhibition of PG formation by ibuprofen. We observed a similar degree of COX inhibition *ex vivo* in both partial and complete responders at this time point; thus, the differences in urinary PG metabolite levels suggest that complete responders had greater activation of the COX pathway *in vivo* in response to surgery. These findings are consistent with prior studies in rheumatoid arthritis and chronic pain patients demonstrating a greater response to NSAID treatment in patients with elevated PGE_2_ levels [44, 45], and support the notion that NSAIDs are most effective in patients in whom activation of the COX pathway is a key mechanism contributing to their pain.

Third molar extraction promoted a systemic inflammatory response, as evidenced by induction of serum cytokines and chemokines and alterations in gene expression in PBMCs, which is consistent with studies in a variety of surgical models [46, 47]. While IL-6, IL-10, and MCP-1 were induced to a similar degree regardless of response group, we observed greater increases in serum TNF-α and IL-8 levels in complete responders compared to partial responders at the second post-surgery time point. Prior studies have shown that PGE_2_ induces both TNF-α [48] and IL-8 [49, 50] *in vitro.* Thus, the induction of these inflammatory mediators in complete responders may be a consequence of greater PGE_2_ formation in response to surgery. However, it is also possible that these pathways are regulated in parallel, and additional studies are necessary to clarify whether there is a causal relationship between activation of the COX pathway and induction of TNF-α and IL-8 *in vivo.* In contrast to some prior studies [51, 52], ibuprofen treatment did not decrease IL-6 levels in our cohort. This may be due to differences in timing of sample collection as our samples were collected approximately 3 and 5 hours after surgery, while effects of NSAIDs on cytokine levels have been observed at later time points (e.g. 12-24 hours after surgery). The results of our gene expression analysis support an anti-inflammatory effect of ibuprofen treatment. Complete responders exhibited much fewer differentially expressed genes after surgery, as well as lower inflammatory gene expression compared to partial responders. Taken together these results suggest that differences in the regulation of the COX pathway and the inflammatory response to surgery contribute to inter-individual variability in the analgesic efficacy of NSAIDs.

There are limitations to our study. Although demographic and clinical characteristics were not statistically different between partial and complete responders in our cohort, we cannot exclude the possibility that these factors contribute to variability in analgesic efficacy of ibuprofen in a larger population. Similarly, our small sample size precluded investigation of genetic variants that might modulate ibuprofen response. Our cohort included only healthy young adults. While this limits potential confounding due to effects of age and comorbidities, we were unable to interrogate the influence of these factors on ibuprofen response. Also, we evaluated only a single dose of ibuprofen over a relatively short sampling period, so we cannot determine the efficacy of repeated dosing or evaluate the effects of ibuprofen on inflammatory mediators or gene expression beyond the acute post-operative period. Finally, our study evaluated ibuprofen response in the setting of acute inflammatory pain, and it is unknown whether similar mechanisms contribute to variation in analgesic efficacy in persistent or chronic pain.

Despite these limitations, our study serves as a foundation for future mechanistic studies to identify predictive biomarkers of NSAID response. In light of the opioid crisis, there is an emphasis on the development of approaches to providing effective pain relief with non-addictive analgesics, including NSAIDs [53]. However, our results, as well as those of prior studies [16, 19, 20], underscore the heterogeneity in the analgesic response to NSAIDs. The ability to prospectively identify patients who would respond to NSAIDs would help limit unnecessary opioid prescriptions in those patients, as well as ensure that patients who would not achieve pain relief with NSAIDs alone have access to additional analgesics.

In conclusion, our results demonstrate that there is marked inter-individual variability in the analgesic efficacy of ibuprofen following third molar extraction surgery that is not explained by differences in clinical characteristics, ibuprofen plasma concentrations, or degree of COX inhibition *ex vivo.* The differences in urinary PG metabolites, serum cytokines, and gene expression in PBMCs suggest that regulation of the inflammatory response to surgery differs between partial and complete responders. Future studies are necessary to elucidate the molecular mechanisms underlying this variability and identify biomarkers that are predictive of ibuprofen response.

## Acknowledgements

This work was partially supported by funds from Pfizer and the Personalized NSAID Therapeutics Consortium (PENTACON: HL117798 (TG)). The Radioimmunoassay and Biomarkers Core at the Diabetes Research Center is supported by NIH DK19525. Dr. FitzGerald is the McNeil Professor of Translational Medicine and Therapeutics.

